# High diversity, abundance and expression of hydrogenases in groundwater

**DOI:** 10.1101/2023.10.03.560699

**Authors:** Shengjie Li, Damon Mosier, Angela Kouris, Pauline Humez, Bernhard Mayer, Marc Strous, Muhe Diao

## Abstract

Hydrogen may be the most important electron donor available in the subsurface. Here we analyze the diversity, abundance and expression of hydrogenases in 5 proteomes, 25 metagenomes and 265 amplicon datasets of groundwaters with diverse geochemistry. A total of 1,772 new [NiFe]-hydrogenase gene sequences were recovered, which almost doubled the number of sequences in a widely used database. [NiFe]-hydrogenases were highly abundant, almost as abundant as the DNA-directed RNA polymerase. The abundance of hydrogenase genes increased with depth from 0 to 129 m. Hydrogenases were present in 502 out of 1,245 metagenome-assembled-genomes. The populations with hydrogenases accounted for ∼50% of all populations. Hydrogenases were actively expressed, making up as much as 5.9% of methanogen proteomes. Most of the newly discovered diversity of hydrogenases was in “Group 3b”, which was linked to sulfur metabolism. “Group 3d” was the most abundant, which was previously linked to fermentation, but we observed this group mainly in methanotrophs and chemoautotrophs. “Group 3a”, associated with methanogenesis, was the most active in proteomes. Two newly discovered groups of [NiFe]-hydrogenases further expanded the biodiversity. Our results highlight the vast diversity, abundance and expression of hydrogenases in the sampled groundwaters, suggesting a high potential for hydrogen oxidation in subsurface habitats.

## Main text

Conversion of water (H_2_O) into hydrogen (H_2_) using solar and wind energy, followed by terawatt-scale storage of hydrogen in the subsurface is currently considered a key aspect of the energy transition [1-3]. One of the potential challenges of this approach is the microbial oxidation of hydrogen which could induce hydrogen loss [4-6]. Our recent work suggested a high potential for microbial hydrogen turnover in groundwaters, based on dissolved hydrogen concentrations, detection of hydrogenotrophic methanogens and isotopic signatures of ^13^C-CH_4_ and ^18^O-O_2_ in 138 samples [7]. Here we focus on diversity and potential functions of hydrogenases by performing metagenomics and proteomics using an expanded sample set encompassing 265 groundwater samples (geochemically characterized and amplicon sequenced, including 25 metagenomes and 5 proteomes) from 138 wells in Alberta (Canada), with sampling depths between 0 and 157 m (Table S2). The groundwaters displayed a range of oxidation states from oxic to completely reduced, accompanied with a wide range of sulfate (>10 g/L to below detection) and methane concentrations (74 mg/L to below detection).

The abundance and expression of different types of hydrogenases were estimated based on unassembled reads, assembled contigs, metagenome-assembled-genomes (MAGs) and proteins in 25 groundwaters. Few [Fe]- and [FeFe]-hydrogenases were present in our data. However, the catalytic subunit of [NiFe]-hydrogenase was almost as abundant as the DNA-directed RNA polymerase (*rpoB*) (Fig. 1a). In 11 out of 25 shotgun-sequenced samples, hydrogenase genes were more abundant than *rpoB* genes, indicating multiple copies of hydrogenase genes per genome. The abundance of hydrogenase/RpoB correlated positively with the sampling depth (*P*=0.017, Fig. 1b). 616 high-quality and 629 medium-quality MAGs were obtained. Hydrogenases were present in 502 MAGs which together accounted on average for 50% of the relative sequence abundance of all MAGs (Fig. 1c, Table S4). In some samples, populations with hydrogenases accounted for >80% of all populations. Although proteomics of groundwater is challenging due to low cell counts, we obtained proteomes of five groundwaters, showing hydrogenases accounted for 0.016%-1.0% of all expressed proteins (Table S5-S9). In proteomes of single populations, the relative abundance of hydrogenases ranged from 0.0026% (Methylomonadaceae) to 5.9% (Methanobacteriaceae) (Fig. 1d). Interestingly, the three methanogens expressed hydrogenases more actively (>1.3%) than the 21 bacterial populations (<0.19%). Thus, hydrogenase genes might be one of the most prevalent genes in the subsurface and active expression indicated these genes were functional.

**Fig. 1.**
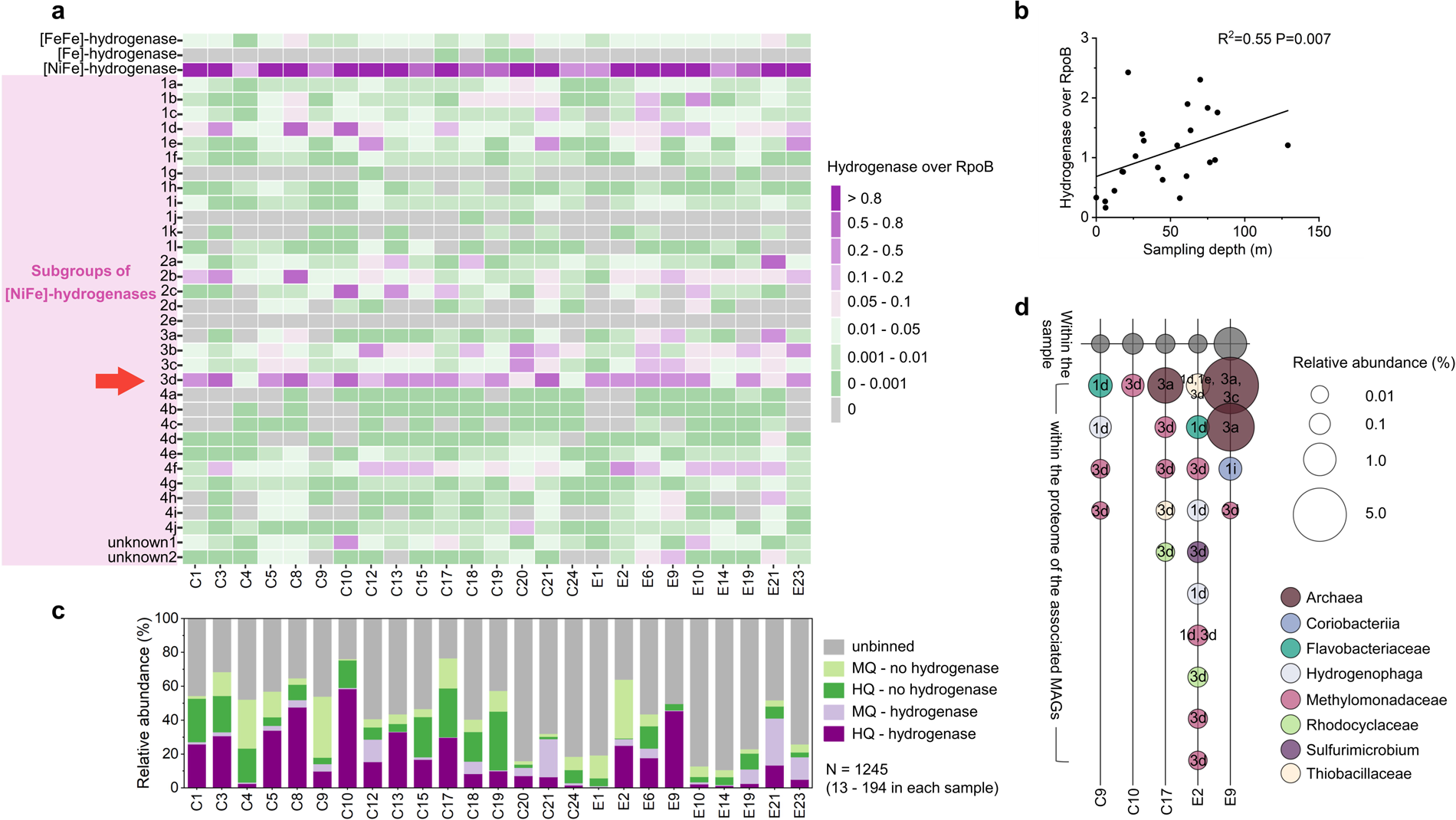
Abundance and expression of hydrogenases. (a) Hydrogenase over RpoB genes calculated from metagenome reads of 25 groundwater samples. The arrow indicates the subgroup of hydrogenases with the highest abundance in sampled groundwaters. (b) Relationship between well sampling depth and total abundance of hydrogenases. Spearman’s rank correlation coefficient and the P value are calculated. The line is the linear regression. (c) Relative abundance of metagenome-assembled-genomes (MAGs) with/without hydrogenases. MQ: medium-quality. HQ: high-quality. There are 1245 MAGs in total. # of MAGs ranges from 13 to 194 in each groundwater sample. (d) Expression of hydrogenases in proteomes. Relative abundance within a sample is calculated as % of all peptide spectral matches of the sample. Relative abundance within a population is calculated as % of all peptide spectral matches of the associated MAG.

From the assembled contigs, 1,772 [NiFe]-hydrogenase gene sequences were recovered, which displayed vast biodiversity (Table S10). These groundwater hydrogenase sequences almost doubled the number (2,014) of [NiFe]-hydrogenase sequences present in a widely used database (Fig. 2) [8]. The newly discovered diversity, abundance and expression were concentrated among a few specific subtypes of [NiFe]-hydrogenases, groups 1e, 3a, 3b, 3d and 4f, with most of the new diversity in 3b, most of the abundance in 3d and most of the expression in 3a.

**Fig. 2.**
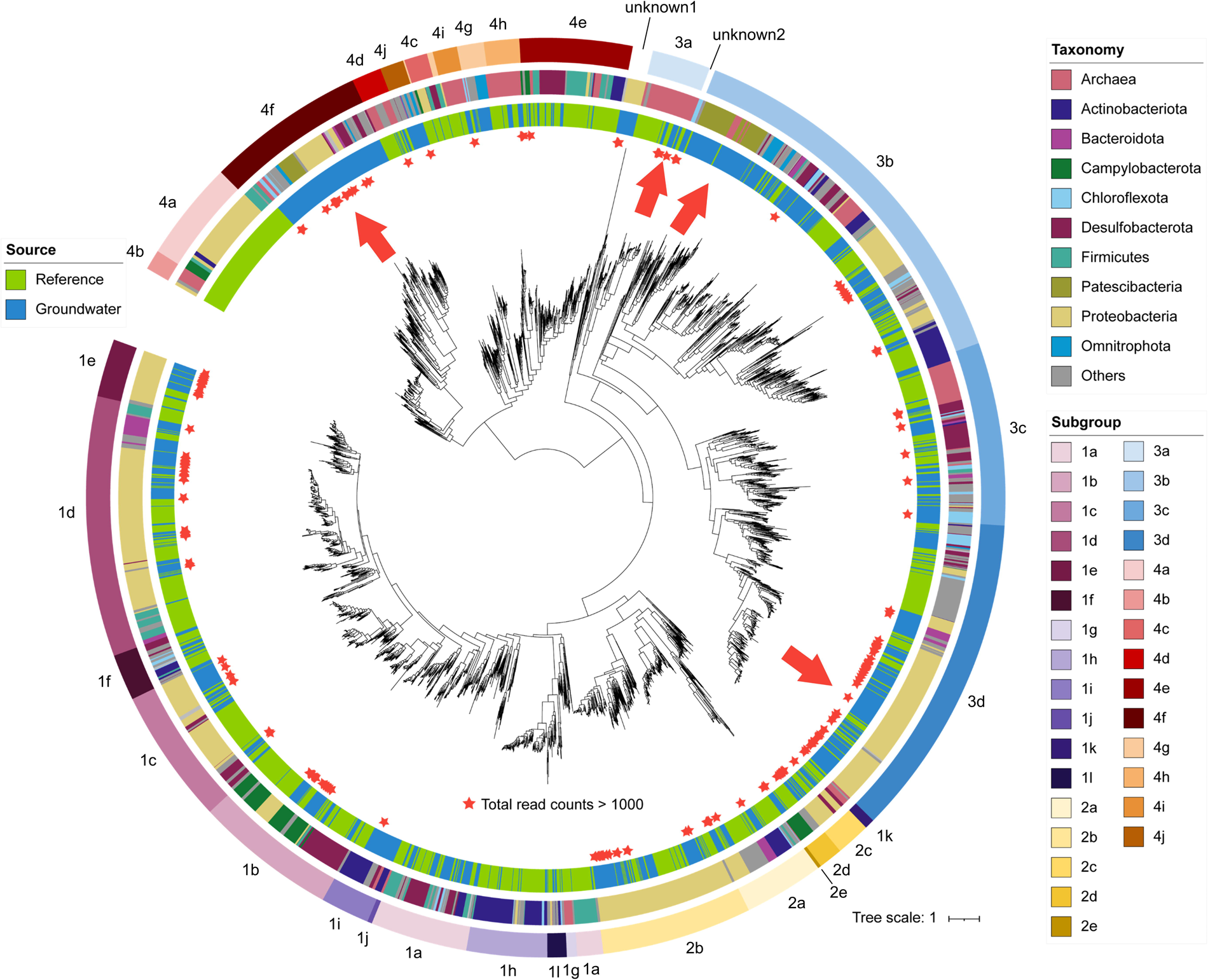
Phylogenetic tree of the catalytic subunit of [NiFe]-hydrogenases. The tree is midpoint-rooted. An arrow inside indicates the place of [NiFe]-hydrogenases with high biodiversity, abundance or expression discovered in sampled groundwaters. Any sequences with total read counts over 1000 in the 25 samples are marked with a star. From inside to outside, the three rings around the tree indicate: (1) source (green, from the reference database; blue, recovered in this study), (2) phylum-level taxonomy and (3) subgroups based on HydDB [5].

Hydrogenases of groups 1e and 3b were previously linked to sulfur reduction [8, 9]. In our data, the abundance of groups 1e and 3b both positively related with sulfate concentration (P=0.005, 0.010, respectively). Based on high-quality MAGs, group 1e was only observed in members of Burkholderiales, and sometimes co-existed with group 3b. Group 3b was typically observed in the genomes of members of Desulfobacterota and Actinobacteriota. For some of these organisms we only detected sulfur oxidation genes, indicating that these hydrogenases might also function alongside sulfur oxidation, coupled to oxygen or nitrate reduction. Many other 3b [NiFe]-hydrogenases were detected in genomes of microbial “dark matter” clades, such as Patescibacteria (14) and Omnitrophota (5), consistent with previous findings [10, 11].

Group 3a was previously proposed to be associated with methanogenesis [8, 9] and was exclusively observed in genomes of methanogens in our data. It was the most highly expressed subgroup in proteomes of MAGs (Fig. 1d). A previous study showed that microbial methanogenesis actively converted CO_2_ to methane in hydrocarbon reservoirs [12]. Both findings indicated high consumption of hydrogen and CO_2_ by methanogens in the subsurface.

Group 3d was previously linked to fermentative metabolism, interconverting electrons between hydrogen and NADH depending on cellular redox state [8, 9]. Group 3d was the most abundant subgroup in 15 out of 25 groundwater samples (Fig. 1a). Surprisingly, 3d hydrogenase genes were present in 89 high-quality MAGs, with 21 of them affiliated with methanotrophic bacteria, Methylomonadaceae. For the other 68 MAGs, 40 of them contained both RuBisCO and phosphoribulokinase, indicating a functional Calvin cycle. Examples include genomes affiliated with Rhodocyclaceae (12), Hydrogenophaga (7), Nitrosomonas (7), Rhodoferax (5). Thus, it is likely that these chemolithoautotrophs can use hydrogen as an additional energy source, with the hydrogenase transferring electrons from H_2_ to NAD^+^ to drive their Calvin cycles.

Two newly discovered groups of [NiFe]-hydrogenases further expanded the biodiversity. The first was positioned near the root of the tree (Fig. 2). This group was composed of 29 sequences, mainly found in Gammaproteobacteria, including the iron oxidizer Sideroxydans, the iron reducer Rhodoferax, the sulfur oxidizer Sulfuritalea and the methanotroph Methylovulum. The other newly discovered group was a sister group of 3a, composed of nine sequences, including four sequences affiliated with archaea and five sequences affiliated with Anaerolineaceae.

Consistency in the types/subgroups of hydrogenases and metabolisms among MAGs with the same taxonomic identity was observed for common groundwater residents, which helped to extrapolate metagenomic results to 265 amplicon-sequenced groundwater samples. For example, total relative abundance of members affiliated with Methylomonadaceae (all 21 MAGs with hydrogenases) could reach 88.6% (Supplementary Table S12). Members of Hydrogenophaga (8 out of 14 MAGs with hydrogenases) could be as abundant as 71.2%. These findings suggest a high potential for hydrogen consumption in sampled subsurface habitats.

Although the subsurface ecosystems analysed here would generally not be considered for hydrogen storage, our study adds to growing evidence that hydrogenases are diverse, functional and ubiquitous in subsurface environments [13-16]. However, with methanogens as the most active hydrogen oxidizers, this need not always be a barrier to hydrogen storage, since recovery of methane could still be a desirable outcome. Likely, any subsurface environment at a temperature conducive to life would harbor resident microorganisms that thrive on hydrogen.

## Data availability

Amplicons in this study are under the Bioproject PRJNA861683 and PRJNA700657. Metagenomes and metagenome-assembled-genomes are under the Bioproject PRJNA700657 (NCBI). The mass spectrometry proteomics data have been deposited to the ProteomeXchange Consortium via the PRIDE partner repository [17] with the dataset identifier PXD044305.

## Acknowledgements

The authors thank the Groundwater Observation Well Network team members of Alberta Environment and Protected Areas (https://www.alberta.ca/groundwater-observation-well-network.aspx) for providing access to groundwater monitoring wells, for sampling and providing highest quality groundwater samples, and for sharing measurement results and expertise. We would like to thank the University of Calgary’s Center for Health Genomics and Informatics for sequencing and informatics services. We thank Carmen Li for help with MiSeq sequencing. We also thank Daan R. Speth for help with phylogenetic analysis. This study was supported by the Natural Sciences and Engineering Research Council (NSERC) through a Discovery Grant and Canada Research Chair (CRC-2020-00257, MS) to Marc Strous, the Canada Foundation for Innovation (CFI), the Digital Research Alliance of Canada, the Canada First Research Excellence Fund (CFREF), the Government of Alberta, and the University of Calgary.

## Author contributions

SL, MS and MD designed the study. SL, DM, AK, PH, BM and MD performed lab research. SL, DM, MS and MD analyzed the data. PH and BM assisted with data interpretation. SL, MS and MD wrote the manuscript.

## Notes

### Competing Interest Statement

The authors have declared no competing interest.

