## Supplementary Information for "High diversity, abundance and expression of hydrogenases in groundwater"

### Groundwater wells and sampling

Each year, approximately 40-90 groundwater wells are monitored and sampled by Groundwater Observation Well Network (GOWN) team members affiliated with Alberta Environment and Protected Areas (https://www.alberta.ca/groundwater-observation-wellnetwork.aspx). For this study, we have collected 265 groundwater samples from 138 wells from 2016 to 2021 (Figure S1). The sampling and physico-chemical measurements of groundwater samples were described in a previous study [1]. Briefly, wells were purged to improve sample quality and samples were collected after field parameters (dissolved oxygen - DO, oxidation reduction potential - ORP, temperature, electrical conductivity - EC) had stabilized. Detailed information of well locations, geology and aqueous geochemistry of the groundwater samples is provided in Table S2.

### DNA extraction and amplicon sequencing

100-1000 mL groundwater samples were filtered through 0.10 μm pore size membrane disc filters (Millipore Sigma). Genomic DNA was extracted from the membrane filter using the DNeasy PowerLyzer PowerSoil Kit (Qiagen, Germany). DNA concentrations were measured with Qubit 2.0 Fluorometer (Thermo Fisher Scientific, Canada).

The V3-V4 region of bacterial 16S rRNA gene was amplified with the primers S-D-Bact-0341a-S-17 (5’-CCTACGGGAGGCAGCAG-3’) and S-D-Bact-0785-a-A-19 (5’GACTACHVGGGTATCTAATCC-3‘) [1, 2]. Polymerase chain reaction (PCR) systems were prepared as previously described [3]. DNA was amplified with the following PCR protocol: an initial denaturation cycle (95°C for 3 min), 25 cycles of denaturation

(95°C for 30 s), annealing (55°C for 45 s) and extension (72°C for 60 s), and a final extension cycle (72 °C for 5 min). Triplicated reactions were conducted for each DNA sample and the PCR products were verified by 1% agarose gel electrophoresis. The amplicons were pooled, purified and sequenced with an Illumina Miseq System (Illumina, San Diego, CA) using the 2 × 300 bp MiSeq Reagent Kit v3. Raw data was processed with amplicon sequencing variant (ASV) analysis in MetaAmp v3.0 [4]. A total of 284 samples, including several technical replicates, yielding 6,559,840 reads after quality control (1,043 to 88,723 reads per sample).

**Metagenomic sequencing, assembly and binning**

DNA of 25 groundwater samples collected in 2019 were selected for metagenomic sequencing. The DNA was fragmented to an average insert size of ~350 bp fragments using a S2 focused-ultrasonicator (Covaris, Woburn, MA). Libraries were prepared using the NEBNext Ultra II DNA Library Prep Kit for Illumina (New England Biolabs, Ipswich, MA) according to the manufacturer's protocol, including size selection with SPRIselect magnetic beads (Beckman Coulter, Indianapolis, IN) and PCR enrichment (eight cycles) with NEBNext Multiplex Oligos for Illumina (New England Biolabs, Ipswich, MA). DNA concentrations were estimated using qPCR and the Kapa Library Quantitation Assay for Illumina (Kapa Biosystems, Wilmington, MA). Genomic DNA was sequenced on an Illumina NovaSeq 600 sequencer (Illumina, San Diego, CA) using a 300 cycle (2 × 150 bp) S1 flow cell. 29,575,481 to 122,086,683 paired reads (4.2 to 17.5 Gb) were generated per sample. Quality trimming of raw reads was performed with BBduk following a previous workflow [3]. Briefly, the last base off of 151 bp reads was trimmed, PhiX sequences were filtered out, adapters and 3’ low quality bases were clipped off.

Trimmed reads of each sample were assembled independently with MEGAHIT v1.2.2-beta [5]. Per contig sequencing depth in reads was determined with BBMap v38.06, with a 95% identity filter. The assembled contigs were binned by three tools, MetaBat v2:2.15 [6], Maxbin v2.2.7 [7] and CONCOCT v1.1.0 [8]. The best metagenome-assembled-genomes (MAGs) obtained from the three binning methods were selected by DASTOOL v1.1.2 [9]. Completeness and contamination of the MAGs were estimated by CheckM2 v0.1.3 [10]. The MAGs were sorted based on the resulting completeness and contamination into high quality (>90% completeness, <5% contamination), medium quality (>50% completeness, <10% contamination) and low quality (<50% completeness, <10% contamination), as is standard according to ENA guidelines (https://ena-docs.readthedocs.io/en/latest/faq/metagenomes.html). The relative abundance of each population associated with a MAG in each metagenome was calculated with the “checkm coverage” and “checkm profile” commands within CheckM v1.1.3b [11]. The taxonomic identity of MAGs was obtained with GTDBtk v2.3.0 [12]. All MAGs and unbinned contigs were annotated using MetaErg v2.3.39 [13]. Only high-quality and medium-quality MAGs were used to check the presence of hydrogenase genes in this study. Only high-quality MAGs were used to link the type/subgroup of hydrogenases and microbial taxonomy in this study.

**Phylogenetic analysis of hydrogenases**

A maximum-likelihood phylogenetic tree of the catalytic subunit of [NiFe]-hydrogenases was built with the following steps. First, all assembled contigs were searched against the reference sequences in the HydDB database [14] with DIAMOND v2.0.9 using a loose criteria "-e 1e-5" [15]. To remove false positives, the hitted sequences were annotated with MetaErg v2.3.39 [13]. Proteins that were not annotated as a hydrogenase or shorter than 150 amino acids were not further considered. Identical sequences were also removed. The remaining 1,772 [NiFe]-hydrogenase sequences from assembled contigs were concatenated with 2,014 [NiFe]-hydrogenase reference sequences in the HydDB. They were aligned with default parameters in MAFFT v7.475 [16]. The phylogenetic tree was constructed with "-m MFP -B 1000" in IQ-TREE v2.0.3 [17] and visualized using iTOL v6.7.4 (https://itol.embl.de/).

**Quantification of hydrogenases in metagenomes**

Two approaches, respectively based on short reads and MAGs, were used to quantify the abundance of hydrogenases in the 25 shotgun-sequenced groundwater samples. For the read-based approach, all quality-controlled reads were searched against the hydrogenase sequences including the references and those retrieved from our samples using DIAMOND v2.0.9 with a loose criteria "-e 1e-5" [15]. To remove false positives, a database was built with all FASTA-formatted protein files in the NCBI RefSeq database [13] for the genomes that were also present in the Genome Taxonomy Database (GTDB) gtdbtk_r207_data.tar (https://data.ace.uq.edu.au/public/gtdb/data/releases/release207/207.0/) [12]. The hitted reads were searched against this large protein database using DIAMOND v2.0.9 [15]. Only the reads that still matched to a hydrogenase sequence were considered. The read counts of hydrogenases were normalized by the read counts of the beta subunit of DNA-directed RNA polymerase, RpoB. For the MAG-based method, any high-quality and medium-quality MAGs with a hydrogenase sequence annotated with MetaErg v2.3.39 [13] were considered.

**Protein extraction and metaproteomics**

Proteins were extracted from 5 groundwater samples. Briefly, 1-20 L samples were filtered through 0.10 µm pore size membrane disc filters (Millipore Sigma). The membranes were cut and transferred to lysing matrix bead tubes A (MP Biomedicals) with the addition of SDT-lysis buffer (0.1M DTT) in a 10:1 ratio [18]. The tubes were bead-beated in an OMNI Bead Ruptor 24 for 45 s at 6 m $s^{-1}$ and then incubated at 95 °C for 10 min. Peptides were isolated from pellets by filter-aided sample preparation (FASP) [19]. Protein concentrations were measured with Qubit 2.0 Fluorometer (Thermo Fisher Scientific, Canada).

Samples were analyzed by 1D-LC-MS/MS. Two wash runs and one blank run were done between samples to reduce carry over. For each run, 2000 ng of peptide solution were loaded onto a 5 mm, 300 µm ID C18 Acclaim® PepMap100 pre-column (Thermo Fisher Scientific) using an UltiMateTM 3000 RSLCnano Liquid Chromatograph (Thermo Fisher Scientific) and desalted on the pre-column. After desalting the peptides, the pre-column was switched in line with a 75 cm × 75 µm analytical EASY-Spray column packed with PepMap RSLC C18, 2 µm material (Thermo Fisher Scientific), which was heated to 60 °C. The analytical column was connected via an Easy-Spray source to a Q Exactive Plus hybrid quadrupole-Orbitrap mass spectrometer (Thermo Fisher Scientific). Peptides were separated on the analytical column using a 460 min gradient as previously described [20] and mass spectra were acquired in the Orbitrap as described previously [21]. 270,275 to 312,887 MS/MS spectra were acquired per sample.

For protein identification, the database of a sample was created using predicted protein sequences of all binned and unbinned contigs from the corresponding metagenome. Proteins with >95% amino acid identity in each database were removed by cd-hit [22], while giving preference to proteins from binned contigs using the "cd-hit-2d" command. The cRAP protein sequence database (http://www.thegpm.org/crap/) containing protein sequences of common laboratory contaminants was appended to the database. The final database of each sample contained 67,020 to 129,602 protein sequences. For protein identification MS/MS spectra were searched against the database using the Sequest HT node in Proteome Discoverer version 2.2.0.388 (Thermo Fisher Scientific, CA, USA) [18]. The Percolator Node and FidoCT were used to estimate false discovery rates (FDR) at the peptide and protein level, respectively. Only proteins identified with medium or high confidence were retained, resulting in an overall false discovery rate of < 5%. [20]. In total, 1,478,976 MS/MS spectra were acquired, yielding 451,848 peptide spectral matches (PSMs) and 34,191proteins of at least "medium" confidence (2,788 to 10,532 proteins per sample). Spectral abundance factor (SAF) of a protein was calculated as the PSM value divided by the number of amino acids. Relative abundance of a protein within a sample was calculated as the SAF value of the protein divided by total SAF value of all proteins. Relative abundance of a protein within a population was calculated as the SAF value of the protein divided by total SAF value of all proteins of the associated MAG.

**Statistical analysis**

The correlation analysis between geochemistry and gene abundance was carried out in PASW Statistics v18.0 using Spearman's rank correlation coefficient. Statistical analyses were considered significant with $P$ values $<0.05$.

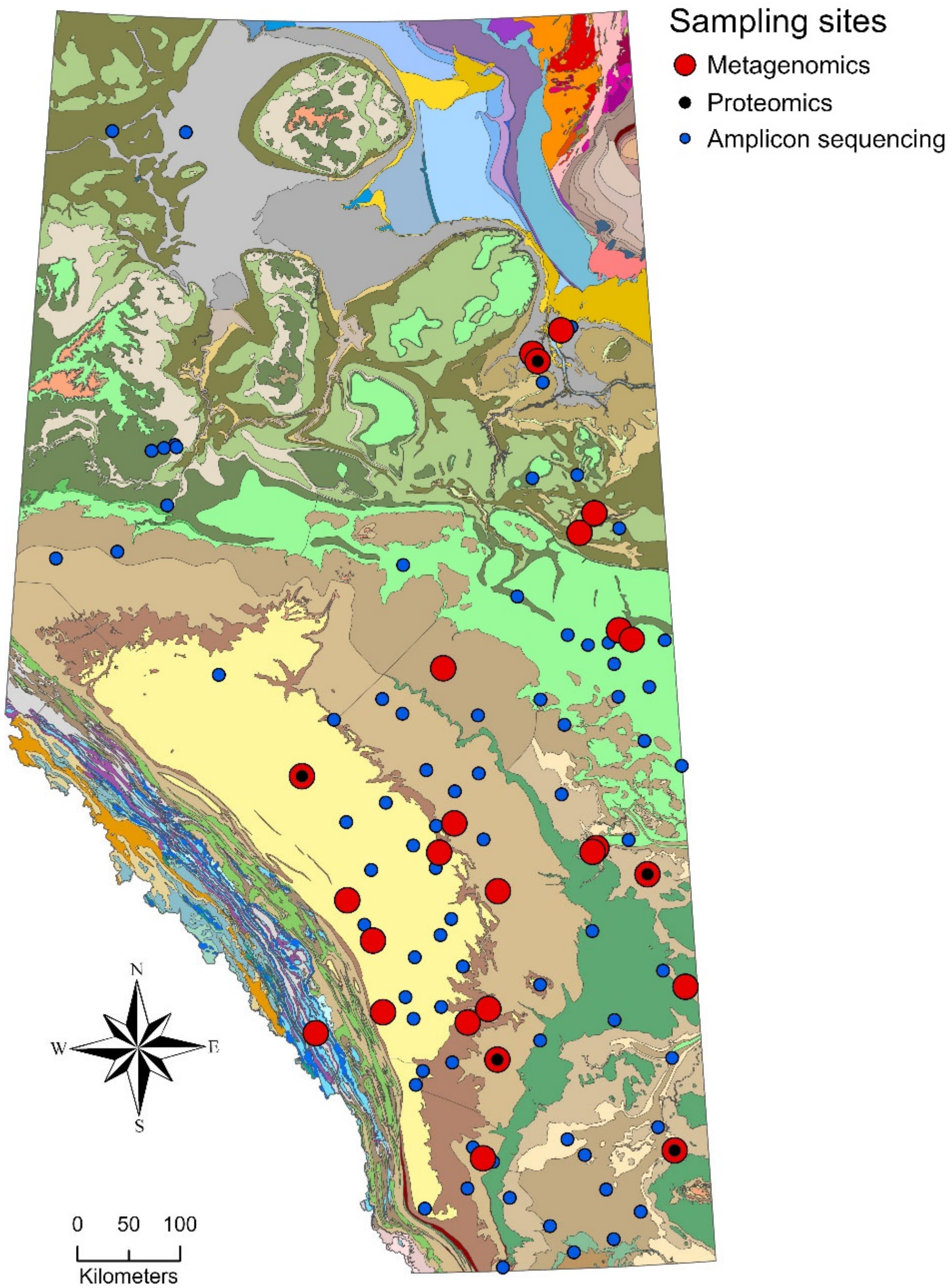


**Fig. S1 Sampling locations of groundwater across Alberta, Canada.** Each circle represents 1 sampling well. Colors indicate the analysis types (red: metagenomics; black: proteomics; blue: amplicon sequencing analysis). The colors in the background indicate bedrock types. The map was created using ArcGIS Pro v2.4.1.
